# NeRFax: An efficient and scalable conversion from the internal representation to Cartesian space

**DOI:** 10.1101/2022.05.25.493427

**Authors:** Oliver Dutton, Falk Hoffmann, Kamil Tamiola

## Abstract

**Motivation:** Accurate modelling of protein ensembles requires sampling of a large number of 3D conformations. A number of sampling approaches that use internal coordinates have been proposed, yet poor performance in the conversion from internal to Cartesian coordinates limits their applicability.

**Results:** We describe here NeRFax, an efficient method for the conversion from internal to Cartesian coordinates that utilizes the platform-agnostic JAX Python library. The relative benefit of NeRFax is demonstrated here, on peptide chain reconstruction tasks. Our novel approach offers 35-175x times performance gains compared to previous state-of-the-art methods, whereas >10,000x speedup is reported in a reconstruction of a biomolecular condensate of 1,000 chains.

**Availability:** NeRFax has purely open-source dependencies and is available at https://github.com/PeptoneLtd/nerfax.

## 1 Introduction

Molecular simulations have proven singular in their capacity to faithfully reproduce complex dynamic behaviour of large biomolecular systems (Hollingsworth and Dror (2018)). They are especially relevant for intrinsically disordered proteins (IDPs), whose highly dynamic ensembles of structures are difficult to study with classical structural biochemistry tools (Uversky and Longhi (2010)). The dynamic behaviour of IDPs defines their ability to form large-weight condensates (Dzuricky *et al*. (2020)), modulate their biological function and renders them prime drugging targets of unmet medical needs (Metallo (2010)). Thus a great effort is devoted to developing theoretical methods for time-efficient sampling of disordered ensembles of arbitrary size and complexity.

Conformational sampling for protein simulations has been previously improved by using an internal coordinates representation for both structural refinement from NMR data (Güntert *et al*. (1997)), molecular dynamics (Chen *et al*. (2005)) and structure generation (Jumper *et al*. (2021)). These involve the transformation from internal to Cartesian coordinates at each step which can be a computational bottleneck.

Parsons *et al*. (2005) introduced the Natural extension of Reference Frame (NeRF) algorithm as the most performant method for Cartesian reconstruction on a singular processor. Subsequently, a parallel variant of NeRF (pNeRF) was developed for reconstruction of unbranched polymer chains (AlQuraishi (2019)). The method was primarily aimed at massively-parallel GPUs. The key limitation of pNeRF was a lack of the placement of protein sidechains. Bayati *et al*. (2020) proposed a refinement of pNeRF that was extended to branched chains. Their method reconstructed a protein in a two-step process that consisted of processing of the longest serial part of the chain, followed by the placement of the branches in completely parallel fashion. The major limitation of the aforementioned approach was poor performance. An arbitrary reconstruction task was taking 2 seconds for a 100 residue protein. Alcaide *et al*. (2022) proposed a generalised implementation of the pNeRF algorithm, denoted mp-NeRF. Their approach utilized the branched chain strategy of Bayati *et al*. (2020) bringing down the reconstruction time to 10 ms on average.

In this work we describe NeRFax, an implementation of NeRF and, to the best of our knowledge, the first fully parallel implementation of pNeRF using the Python JAX library. The relative benefit of our approach is demonstrated for arbitrary single chain proteins and a medically relevant biomolecular condensate.

## 2 Approach and Implementation

### 2.1 Framework

Unlike a number of previous implementations of NeRF that leveraged C++ - like performance and accessibility of the high-level Python libraries Tensorflow and Pytorch (Alcaide *et al*. (2022); AlQuraishi (2019)), our algorithm was programmed using JAX. Superior numerical performance of NeRFax transpires from optimisations applied during compilation, including fusion and constant folding. An immediate consequence of these optimisations is increase in hardware utilisation through reduced memory transfer and a simplified computational graph due to pre-calculation of values known at compilation. The automatic vectorisation capability of JAX, owing to its functional nature, accelerates development so that code for reconstructing multiple chains in parallel was generated by the addition of just one extra line of code to the single chain variant (Frostig *et al*. (2018)).

### 2.2 Parallelisation

The initial pNeRF implementation by AlQuraishi (2019) focused solely on the reconstruction of the protein backbone, an unbranched chain of -N-CA-C- repeating units. The parallelisation in pNeRF was achieved by splitting the protein into *M* fragments, each of ≥ 3 particles, which were reconstructed in parallel in an arbitrary reference frame and concatenated in a sequential manner. As a consequence, for a system of *N* atoms, there were *N/M* placement steps and *M* stitching steps, which led to linearly increasing execution time. pNeRF method was further improved by changing the number of protein fragments to *M* = *N/*3, as each concatenation step is faster than placement, yielding an *O*(*N*) performance bottleneck in concatenation process (Alcaide *et al*. (2022)). The concatenation procedure had two *O*(*N*) operations given by the transformation in Equation 1, referred to as an iterative matrix-matrix multiplication and a cumulative sum. Here, *a*_0_ ⊕ *a*_1_ represents either a matrix-matrix multiplication (*a*_1_ · *a*_0_) or an addition (*a*_0_ + *a*_1_) operation.

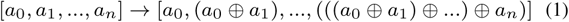

We discovered that as both operators, ⊕, are associative, i.e. (*a* ⊕ *b*) ⊕*c* ≡ *a* ⊕ (*b*⊕ *c*), they can be parallelised by a binary tree scheme illustrated in Supporting Information which reduces the calculation from *O*(*N*) to *O*(log *N*) steps for highly parallel architectures (Hillis and Steele (1986); Blelloch (1990)). Consequently this implementation scales *O*(log *N*) in comparison to *O*(*N*) in previous works.

## 3 Results and Conclusions

A single-CPU implementation of our algorithm, NeRFax, consistently outperformed the state-of-the art NeRF code for every tested protein chain length in a range of 10 to 1,000 residues yielding 35 to 175 speedup (Figure 1). Subsequently, we decided to test our NeRFax method for a biomolecular condensate which consisted of 1,000 protein chains and 1,212,000 atoms. We selected the low complexity domain (LCD) of the RNA-binding protein fused in sarcoma (FUS), as the formation of its biomolecular condensate was recently demonstrated with coarse-grained molecular dynamics simulations (Benayad *et al*. (2021)). Our NeRFax method performed a full-scale reconstruction in 3.4 ms on a singular A100 GPU, approximately 17,000x faster than the forcibly serial usage of any previous implementation. Specifics of the benchmark are detailed in Supporting Information.

**Fig. 1.**
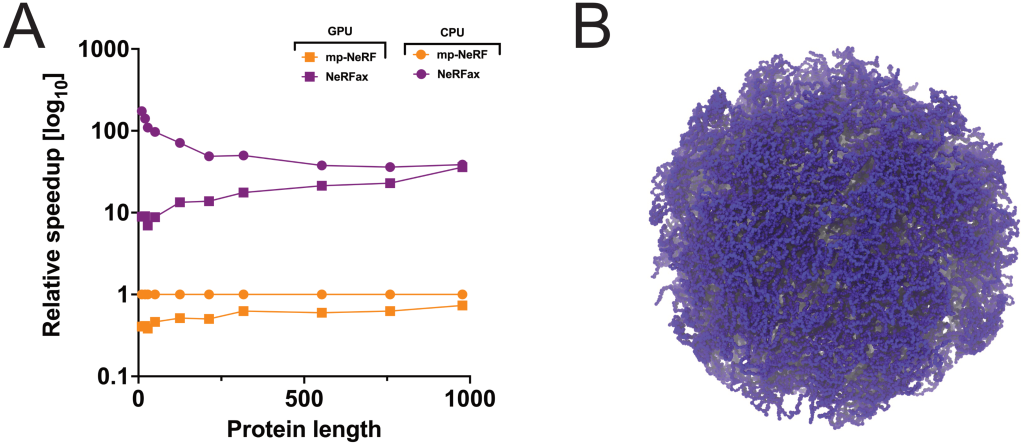
A comparative assessment of performance gain through NeRFax method. The PyTorch mp-NeRF implementation is used as a performance benchmark; A) relative performance on CPU and GPU with respect to the state-of-the-art NeRF code (Alcaide et al. (2022)) as a function of protein length, B) a 1,000-molecule model of an IDP aggregate.

We hope that consistent performance gains offered by NeRFax will propel a development of numerically efficient methods for protein structure reconstruction especially for highly dynamic and readily aggregating intrinsically disordered proteins of unmet medical needs.

## Supplementary information

Supplementary data are available at *Bioinformatics* online.

