## Supporting Information for "NeRFax: An efficient and scalable conversion from the internal representation to Cartesian space"

### NeRFax: Supporting Information

#### 1 Parallelisation details

To illustrate how associative cumulative computations can be parallelised we show an example of a cumulative sum of four elements. Figure S1 shows a serial cumulative sum taking three steps. Figure S2 shows a scheme which on hardware able to compute two operations in parallel takes just two steps.

For associative cumulative computations of more than four elements, there are two variants of the parallelisation scheme. The first is work-efficient, taking $2\log_2 N - 2$ steps and $O(N)$ operations. The second is work-inefficient taking just $\log_2 N$ steps but involves $O(N \log N)$ operations.

#### 2 Benchmarking

##### 2.1 Hardware

All the calculations were performed on NVIDIA DGX A100 supercomputer node equipped with AMD EPYC 7742 64C 2.25GHz processors and A100 40GB GPUs.

##### 2.2 Implementation

We implemented and benchmarked three variants of NeRF with varying level of parallelism in protein backbone reconstruction. Our implementations are a serial variant of NeRF method with maximum vectorisation, and two maximally parallelised versions of the mp-NeRF algorithm with work-efficient and work-inefficient parallelism respectively. Only the most performant method is reported for each case.

In the case of single chain construction on CPU the serial variant is fastest, due to the low parallelism of CPUs. For GPU, the work-inefficient parallelism is the most performant for single chain. While when reconstructing the 1,000 chain condensate the work-efficient is faster, indicating the GPU parallelism is not saturated by 1,000 chains in parallel with the serial scheme, but oversaturated by the work-inefficient scheme.

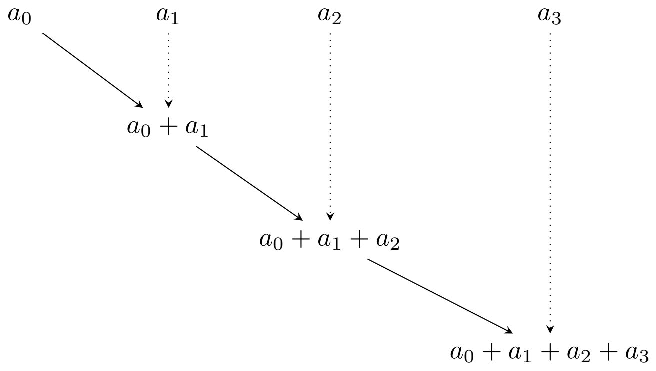


Figure S1: Serial implementation of cumulative sum of four elements, taking three steps and three operations

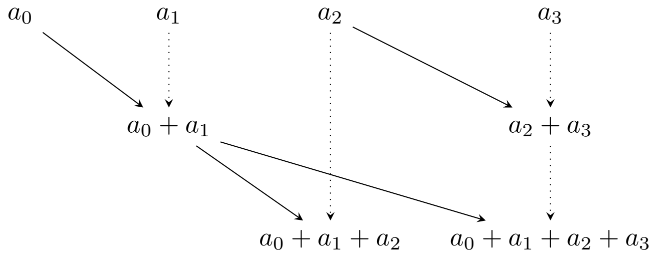


Figure S2: Parallel implementation of cumulative sum of four elements, taking two steps and four operations
